# Polishing Copy Number Variant Calls on Exome Sequencing Data via Deep Learning

**DOI:** 10.1101/2020.05.09.086082

**Authors:** Furkan Özden, Can Alkan, A. Ercüment Çiçek

## Abstract

Accurate and efficient detection of copy number variants (CNVs) is of critical importance due to their significant association with complex genetic diseases. Although algorithms that use whole genome sequencing (WGS) data provide stable results with mostly-valid statistical assumptions, copy number detection on whole exome sequencing (WES) data shows comparatively lower accuracy. This is unfortunate as WES data is cost efficient, compact and is relatively ubiquitous. The bottleneck is primarily due to non-contiguous nature of the targeted capture: biases in targeted genomic hybridization, GC content, targeting probes, and sample batching during sequencing. Here, we present a novel deep learning model, *DECoNT*, which uses the matched WES and WGS data and learns to correct the copy number variations reported by any off-the-shelf WES-based germline CNV caller. We train DECoNT on the 1000 Genomes Project data, and we show that we can efficiently triple the duplication call precision and double the deletion call precision of the state-of-the-art algorithms. We also show that our model consistently improves the performance independent from (i) sequencing technology, (ii) exome capture kit and (iii) CNV caller. Using DECoNT as a universal exome CNV call polisher has the potential to improve the reliability of germline CNV detection on WES data sets.

## 1 Introduction

Gene copy number polymorphism in a population due to deletions and duplications of genomic segments substantially derive genetic diversity [31,15], affecting roughly 7% of the genome [34]. This class of structural variations (SVs), called copy number variations (CNVs), have also been associated with several genetic diseases and disorders such as neurodevelopmental/neurodegenerative disorders [27,11,24,7,43] and various cancers such as breast, ovary, and pancreas cancers [12,23,29,26]. Karyotyping and microarray analyses have been the standard clinical testing for disease-causing CNVs for many years [40], but High Throughput Sequencing (HTS) has all but replaced these techniques with the ability to theoretically capture all forms of genomic variation. Numerous CNV detection algorithms have enjoyed success by analyzing whole genome sequencing (WGS) data using different sequence signatures such as read depth, discordant paired-end read mappings, and split reads [13]. WGS is a convenient resource for CNV callers as it provides near-Poisson depth of coverage [3]. On the other hand, accurate CNV detection on whole exome sequencing (WES) data has mostly been lacking. The algorithms which call CNVs on the WES data have notoriously high false discovery rates (FDR) reaching up to ∼60% which renders them impractical for clinical use [42,36]. This is mainly due to several problems associated with the WES technology such as non-uniform read-depth distribution among exons caused by biases in (i) sample batches, (ii) GC content, and (iii) targeting probes [17,22,19]. This is unfortunate as WES data size is ten times smaller (i.e., ∼10GB vs ∼100GB) and it costs three times less compared to WGS which makes it highly abundant and a common choice for analyzing complex genetic disorders [33,8,30,32]. For instance, the Genome Aggregation Database (gnomAD) contains around 125K WES samples as opposed to 70K WGS samples [18]. Thus, currently, such a rich resource of large scale WES data cannot be fully utilized to investigate the contribution of copy number variation to disease etiology.

Here, we present the first of its kind, exome CNV call *polisher* named *DECoNT* (Deep Exome Copy Number Tuner) to improve the performance of any off-the-shelf WES-based germline CNV detection algorithm. DECoNT is a deep learner that utilizes matched WES and WGS samples present in the 1000 Genomes Project [39] data set to learn the association between (i) calls made by any CNV caller that use WES data and (ii) ground truth calls generated from the WGS data for the same sample. Based on a bidirectional long short-term memory (Bi-LSTM) based architecture, it uses only WES read depth along with the calls from a third party caller and learns to correct noisy predictions (Figure 1). We show that DECoNT can improve the duplication and deletion call precision of the state-of-the-art algorithms by up to 3-fold and 2-fold, respectively. The performance gain is consistent among CNV callers that output integer copy number predictions and categorical predictions (i.e., deletion, duplication, or no call). As the training phase is offline, polishing procedure is memory and time efficient and takes only a few seconds on average per sample. Furthermore, we show that the models learned are universal in the sense that they are (i) sequencing platform, (ii) target capture kit, and (iii) CNV caller independent. For instance, using models learned on 1000 Genomes Project data set that uses Illumina as the sequencing platform and various capture kits such as Agilent and NimbleGen, DECoNT can correct calls made by any of the state-of-the-art CNV caller, for samples obtained from other capture kits like Agilent SureSelect and Illumina Nextera Exome Enrichment Kit or different sequencing platforms like Illumina HiSeq 4000, Illumina NovaSeq 6000, and MGI; that are “unseen” during training. Thus, DECoNT is highly flexible and scalable and makes exome based CNV detection practical by boosting the performance of virtually any WES-based CNV caller algorithm. The tool and the models are available at https://github.com/ciceklab/DECoNT. All necessary scripts and data to replicate the results presented in the figures and tables are deposited to https://zenodo.org/record/3865380.

**Fig. 1.**
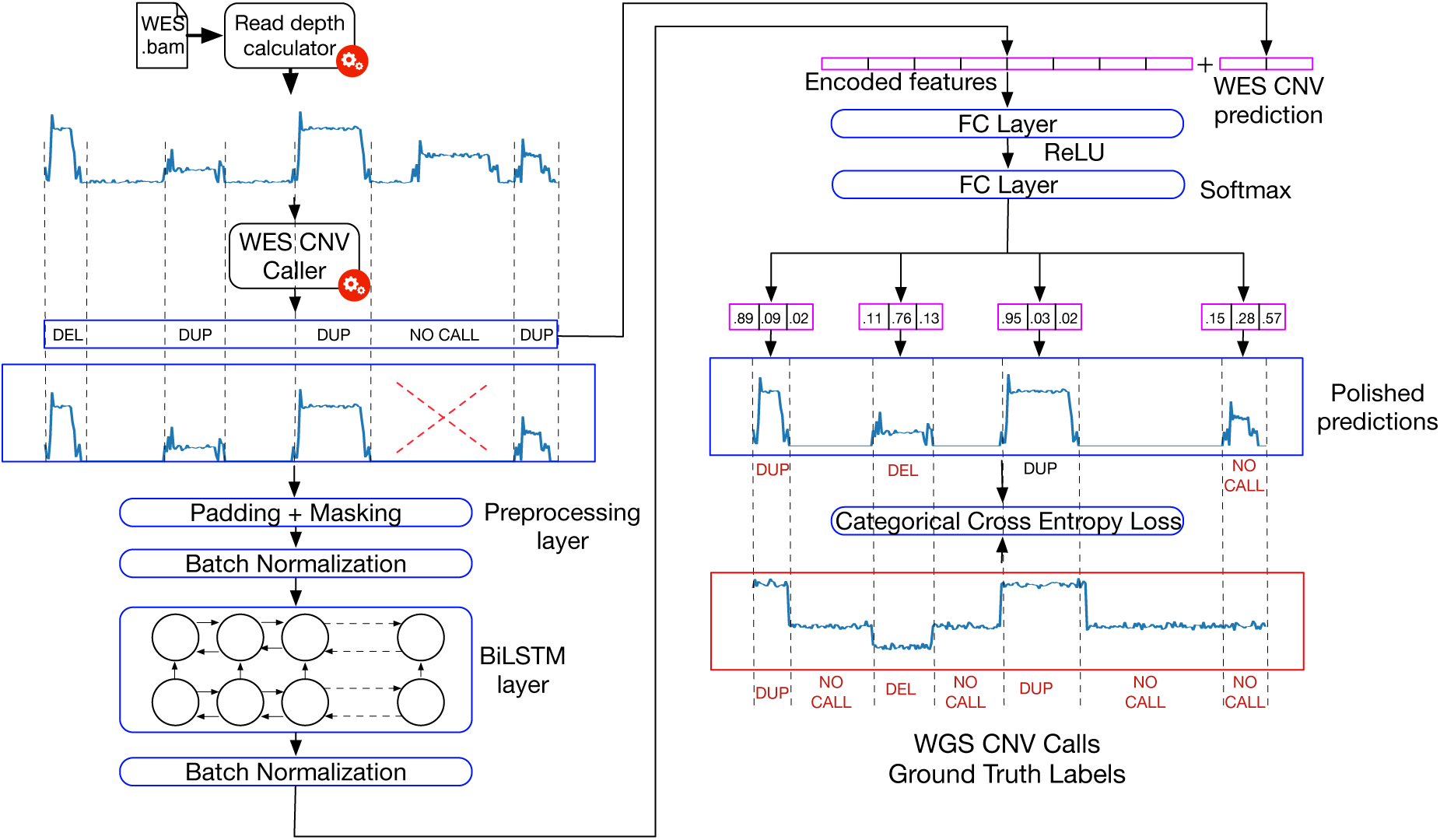
Learning workflow of DECoNT. First, BAM file that corresponds to a WES data set from **1000 Genomes data set** is used to calculate exome-wide read depth which is input to a third party WES-based CNV caller. The caller generates the calls for various regions which could be (i) a binary prediction like duplication, deletion (e.g., XHMM [9]) as shown in the figure, or (ii) an integer value that indicates the exact copy number (i.e., Control-FREEC [4]). The read depth of the regions for which a call has been made is input to a Bi-LSTM model. Encoded features are passed from a series of fully connected layers along with the original prediction of the caller algorithm. Using the ground truth calls from the WGS data of the same sample the method learns to predict (correct) the calls using cross entropy loss for the binary outputs (as shown in the figure) and using mean squared loss for integral calls.

## 2 Results

### 2.1 Bi-LSTM based Neural Network Learns to Correct False Positive Germline WES CNV Calls

A Bidirectional Long Short-Term Memory (Bi-LSTM) network [14] is a type of recurrent neural network which learns a representation (i.e., embedding) of a sequence by processing it for each time-step (i.e, each read depth value in the CNV region in our case) in the forward and the backward directions. While doing so, it remembers a summary of the sequence observed so far to capture the context for each time-step. RNNs and LSTM-based architectures have been widely and successfully used in natural language processing domain to process sequence data [41].

DECoNT uses a single hidden layered Bi-LSTM architecture with 128 hidden neurons in each direction to process the read depth signal (Methods). First, WES-based germline CNV caller result is obtained along with the read depth signal in those event regions. Bi-LSTM subnetwork learns a transformed representation for the read depth sequence (Figure 1). This embedding and the corresponding CNV call are input to a fully connected (FC) layer feed forward neural network. The FC layers predict the polished result for call. DECoNT makes use of the calls made on the WGS data of the same sample as the ground truth for the learning procedure. We use matched WGS data to obtain the ground truth calls for the CNV events called on the WES samples of the same individuals in the 1000 Genomes data set [39].

We polish state-of-the-art WES-based germline CNV callers. There are two types of such algorithms. The first type makes discrete predictions for CNVs (i.e., deletion and duplication). We consider three methods in this category: (i) XHMM [9], (ii) CoNIFER [21], and (iii) CODEX2 [16]. The second type predicts the exact copy number as an integer value. The examples we consider of this type is Control-FREEC [4] and CNVkit [35]. DECoNT architecture is flexible and can be easily modified to polish both types of algorithms (Methods). We train a DECoNT model for every above-mentioned tool using 3 NVIDIA GeForce RTX 2080 Ti and 1 NVIDIA TITAN RTX GPUs in parallel with training times ranging from ∼ 1 to ∼ 4 days (Methods).

We find that DECoNT is able to substantially improve the performance of all algorithms in almost all comparisons. For algorithms that make discrete predictions, we observe improvements in both duplication and deletion call precision (Figure 2a). The largest gain is in duplication call precision for CoNIFER which is improved by 3-fold (i.e., 24.68% to 75%). The largest gain in deletion call precision is again obtained for CoNIFER, which is improved by 1.5-fold (i.e., 45.45% to 68.51%). Also, the overall precision is improved by 2.6-fold (i.e., 27.22% to 71.11%) for CoNIFER. This improvement is especially striking as CoNIFER is relatively conservative compared to other algorithms and seldom make calls despite relaxation of its parameters. For XHMM, we observe 1.4, 1.7, and 1.5 fold increases which correspond to 20%, 29%, and 24% improvements in duplication, deletion and overall precision, respectively. We see a similar trend for CODEX2. Before polishing with DECoNT, CODEX2 achieves 12% duplication precision, 45% deletion precision, and 27% overall precision. DECoNT provides 1.9 fold increase in duplication call precision, 1.5 fold increase in deletion call precision, and 1.75 fold increase in overall precision, respectively. These correspond to 11%, 23%, and 20% improvements in each respective metric. Confusion matrices before and after polishing are shown in Supplementary Figure 1 for all tools. We would like to note that these improvements are obtained in seconds per sample at the test time. Increased precision is an important result for the life scientists who work with these calls as the reliability of the calls are substantially increased as the number of false positives are substantially decreased.

**Fig. 2.**
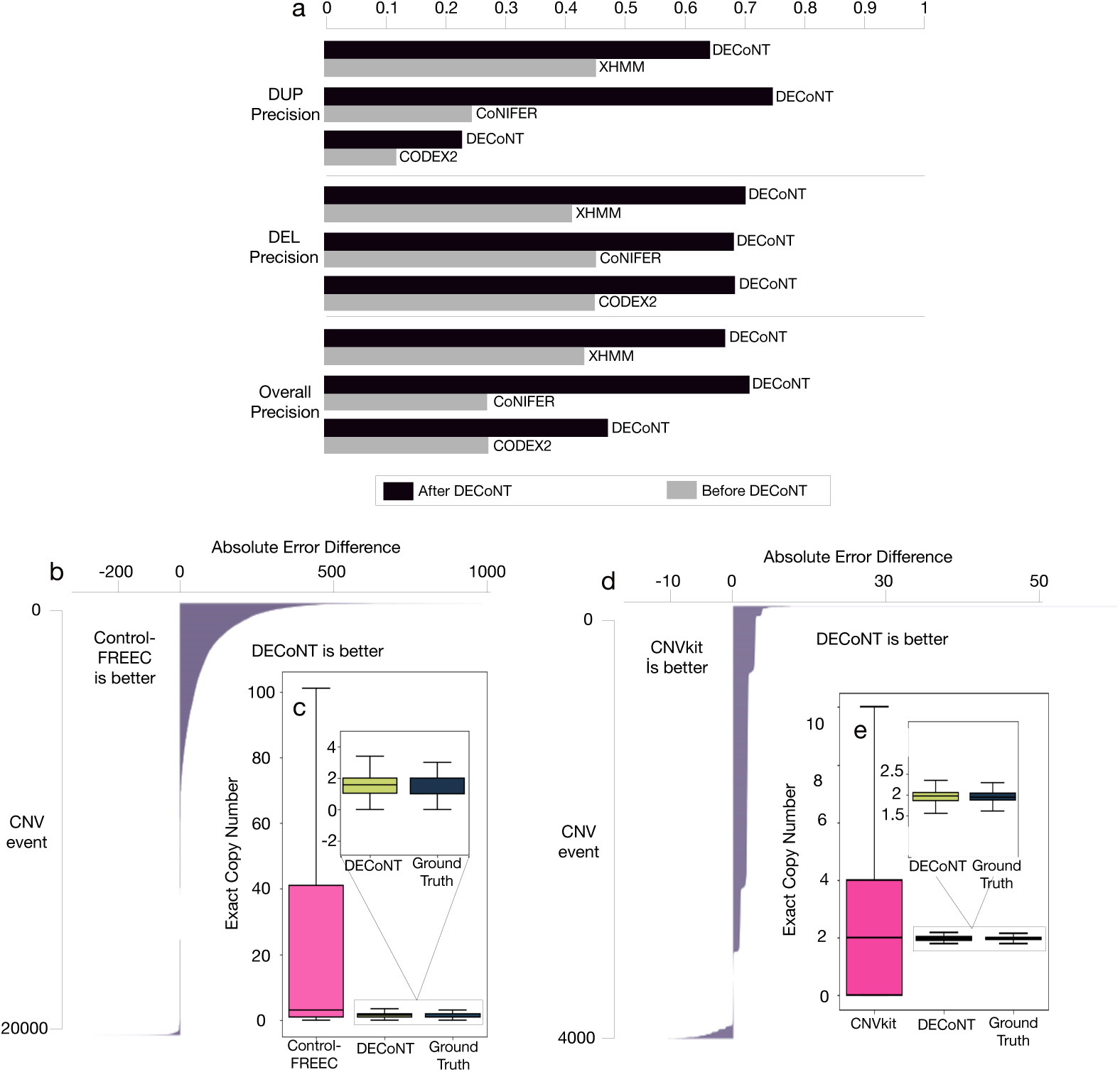
The performance comparison of the WES-based CNV caller’s before and after polishing with DECoNT. a) For the tools which predict existence of a CNV event (XHMM, CoNIFER and CODEX2) are evaluated with respect to duplication call precision, deletion call precision and overall precision. DECoNT improves the performance for all tools in all settings and results in drastic improvements. Different shades of gray represent different tools and the attached black bars represent the DECoNT-polished version of those tools. b) In this panel, we compare Control-FREEC and the DECoNT-polished with respect to Absolute Error (AE) difference on each sample (i.e., events). Bars to the right indicate the magnitude of the improvement due to polishing of DECoNT. For more than half of the samples, DECoNT results show improvement. c) The distribution of the unpolished Control-FREEC predictions in the test samples (pink) is quite different than the ground truth copy number variation distribution. On the other hand, DECoNT polished versions of the same events (dark blue) highly resemble the distribution of the ground truth calls. Black lines across the boxes are median lines for the distributions. Black vertical lines are whiskers and 1.5*×* Inter Quartile Range is defined with the horizontal lines at the top and bottom of the whiskers. 1000 Genomes data set WGS samples are used as ground truth calls in all analyses. Panels d) and e) show the results for CNVkit, similar to b) and c).

As for the Control-FREEC and CNVkit, which output exact copy number values, we evaluate their performance (i.e., absolute error; AE). For Control-FREEC we consider 20, 482 CNV events called (Figure 2b, Methods). DECoNT improves the absolute error in 74.58% of the test samples for an average *AE* improvement of 47.39 and deteriorates the performance in 25.35% of the test samples for an average *AE* deterioration of only 1.2. While unpolished Control-FREEC predictions have a Spearman correlation coefficient of 0.227 with matched ground truth copy numbers, DECoNT-polished predictions have a Spearman correlation coefficient of 0.568 (Figure 2c). DECoNT-polished predictions highly resemble the distribution of the ground truth calls. In order to mimic the discrete prediction case, we also discretized the CNV calls of Control-FREEC to Deletion (CN *<* 2), Duplication (CN *>* 2) and No-Call (CN = 2) categories to measure precision as defined in Section 4.3. Again, DECoNT was able to improve the DEL and DUP precision up to 3-fold (see Supplementary Note 1 for details). The other tool that outputs exact copy number values is CNVkit. We evaluate its performance on 3, 972 CNV events called (Figure 2d, Methods). DECoNT improves the absolute error in 86.78% of the test samples for an average *AE* improvement of 1.82 and deteriorates the performance in 13.21% of the test samples for an average *AE* deterioration of only 0.66. Raw CNVkit predictions have a Spearman correlation coefficient of 0.0156 and DECoNT-polished predictions have a Spearman correlation coefficient of 0.122 (Figure 2e). Similar to Control-FREEC, DECoNT-polished CNVkit predictions highly resemble the distribution of the ground truth calls.

To show the need for a complex model like DECoNT in this application, we used standard machine learning algorithms for polishing and compared the performance. We used SVM and logistic regression for the discrete prediction case and polynomial regression for the exact prediction (rounded). We show that these models actually deteriorate the baseline caller performance and we need more complex models like DECoNT for this task. Details are given in Supplementary Note 2.

We also investigated DECoNT’s polishing performance on a consensus WES-based germline CNV caller, CNLearn [28]. CNLearn first runs 4 WES-based callers (CANOES, CODEX, CLAMMS, XHMM) and then using a random forest classifier, learns to aggregate the results of these programs. We obtained the 39 CNV predictions of CNLearn on 4 samples from the 1000 Genomes Project (personal communication with S. Girirajan; default settings are used). The list of these samples are given in Supplementary Note 4. Using the CNVnator calls obtained on the WGS data of the samples as the ground truth, we observed that CNLearn achieved a precision of 0.79. Using the DECoNT model trained using XHMM calls, we polished the results of CNLearn and improved the precision to 0.889. Note that CNLearn requires more computation as it employs many models, yet, DECoNT was able to improve the performance, even when using a cross-model polisher. More specifically, CNLearn and polished-CNLearn do not agree on 8 calls, out of which, polished version is correct in 4, unpolished version is correct in 2 and both are incorrect in 2. The list of these calls is also given in Supplementary Note 5.

### 2.2 Polishing Performance on a Validated CNV call set

In order to further test the polishing performance of DECoNT, we also use a highly validated CNV call set published by Chaisson et. al., [6]. This data set contains the WGS-based CNV calls of 9 individuals selected from the 1000 Genomes Project samples, for which a consensus call set is obtained using 15 different WGS-based CNV callers with comparisons against high quality SVs generated using long read Pacific Biosciences data with a single basepair breakpoint resolution (Methods). We use data from 8 samples with matched WES data set.

Using the same models explained in Section 2.1, we correct the CNV calls made on WES data of the 8 samples made by XHMM, CoNIFER and CODEX2. Note that none of the DECoNT models have seen the data of these individuals during training. We validate the performance using this call set. Table 1 summarizes the performances before and after polishing with DECoNT, with respect to WGS validated calls.

**Table 1.** The performances of the WES-based CNV caller algorithms before and after polishing are shown (DEL, DUP and overall precision).

| Tool | DUP Precision |  | DEL Precision |  | Overall Precision |  |
| --- | --- | --- | --- | --- | --- | --- |
|  | default | polished | default | polished | default | polished |
| XHMM | 0.064 | **0.071** | 0.257 | **0.387** | 0.135 | **0.170** |
| CoNIFER | 0.090 | **0.160** | 0.0* | **0.314** | 0.090 | **0.250** |
| CODEX2 | 0.027 | **0.046** | 0.387 | **0.685** | 0.185 | **0.350** |

ValidatedWGS CNV call set of Chaisson *et al.* [6] is used as the ground truth CNV call set. We first use matched WES reads to call WES CNVs using CoNIFER, CODEX2, and XHMM. Then, we use DECoNT to polish obtained CNV calls. Table shows the DEL, DUP and overall precision of the methods. \*CoNIFER does not report any deletion events on this set of WES samples.

Similar to analysis above, DECoNT improves the performance of all three algorithms in all comparisons. The most substantial improvements are observed for CoNIFER. 7%, 31.4% and 16% improvements are observed for duplication, deletion, and overall precision, respectively. It is noteworthy that while CoNIFER does not report any deletion events, DECoNT was able to correct incorrect duplication calls into correct deletion calls and increase the precision to 31.4% in this category. See Supplementary Figure 2 for the confusion matrices obtained before and after polishing by DECoNT. For XHMM and CODEX2, we see consistent improvements reaching up to nearly 2-fold for CODEX2.

### 2.3 Polishing Performance Generalizes to Unseen Sequencing Platforms

We obtained the training data from the 1000 Genomes data set, in which the WES component is produced using Illumina Genome Analyzer II and Illumina HiSeq 2000. While data from these platforms is abundant and sufficient training data set size can be met, for users using other sequencing platforms, it might not be possible to train DECoNT due to lack of matched WES and WGS samples. We therefore evaluated whether models trained on the available 1000 Genomes data can be used to polish CNV calls made on WES samples obtained using other sequencing platforms or capture kits that have not been seen by DECoNT (Methods).

We obtain the WES data for the sample NA12878, sequenced using four different platforms: (i) Illumina NovaSeq 6000; (ii) Illumina HiSeq 4000; (iii) BGISEQ-500; and (iv) MGISEQ-2000. We use these four samples only for testing. All considered WES-based CNV callers are used to call CNV events on these four WES samples.

Even though DECoNT has not seen the read depth information or the CNV events on these sequencing platforms, it still can generalize from the training on the 1000 Genomes data and still can substantially improve the performances of XHMM, CoNIFER, and CODEX2 (Table 2 and Supplementary Figure 3). We observe improvements in 34 out of 36 tests.

**Table 2.** The performance of discrete germline WES-based CNV callers on NA12878 data before and after being polished by DECoNT.

| Platform | Tool | DUP Precision |  | DEL Precision |  | Overall Precision |  |
| --- | --- | --- | --- | --- | --- | --- | --- |
|  |  | default | polished | default | polished | default | polished |
| NovaSeq 6000 | XHMM | **0.660** | 0.330 | 0.078 | **0.111** | 0.097 | **0.133** |
|  | CoNIFER | NA* | NA* | NA* | NA* | NA* | NA* |
|  | CODEX2 | 0.043 | **0.139** | 0.198 | **0.398** | 0.112 | **0.266** |
| HiSeq 4000 | XHMM | **0.500** | 0.125 | 0.093 | **0.156** | 0.100 | **0.152** |
|  | CoNIFER | 0.0** | **0.500** | 0.191 | **0.192** | 0.191 | **0.214** |
|  | CODEX2 | 0.032 | **0.075** | 0.188 | **0.389** | 0.099 | **0.212** |
| BGISEQ-500 | XHMM | 0.045 | **0.076** | 0.157 | **0.176** | 0.088 | **0.200** |
|  | CoNIFER | 0.0** | **0.010** | 0.052 | **0.082** | 0.052 | **0.055** |
|  | CODEX2 | 0.051 | **0.156** | 0.214 | **0.492** | 0.125 | **0.364** |
| MGISEQ-2000 | XHMM | 0.045 | **0.076** | 0.157 | **0.176** | 0.088 | **0.200** |
|  | CoNIFER | 0.0** | **0.010** | 0.052 | **0.082** | 0.052 | **0.055** |
|  | CODEX2 | 0.051 | **0.156** | 0.214 | **0.492** | 0.125 | **0.364** |

The most substantial improvement is observed for CODEX2 that corresponds to an average 2.6-fold increase in performance. This even exceeds testing performance on the same platform as training (i.e., ∼ 2-fold improvement). For XHMM, the performance is improved 10 out of 12 tests, reaching up to doubling the performance in overall precision performance for BGISEQ and MGISEQ platforms. For NovaSeq 6000 and HiSeq 4000, the performance deteriorates in duplication precision. However, XHMM makes a few duplication calls: 3 and 2, respectively. While DECoNT keeps the true positives, it adds a few false positives and this results in the performance decrease in these settings. CoNIFER does not report any events on the NovaSeq 6000 platform despite tuning its parameters to more relaxed settings. On BGISEQ-500 and MGISEQ-2000 platforms, even though CoNIFER does not report any duplication calls, DECoNT finds some deletion calls and increases duplication precision from 0 to 1%. While it does not report ant duplication calls for the HiSeq 4000 platform, DECoNT is able to increase the precision to 50% but by reporting a true positive and a false positive. The trend in deletion precision performance is similar. Finally, overall precision performance is consistently increased in all tests and the improvement ranges from 0.3% to 2.3%.

For Control-FREEC, ∼65% to ∼74% of the CNV calls have been improved as opposed to only ∼7% to ∼8% of the calls have been deteriorated by DECoNT. We observe a decrease in average absolute error after polishing in all four platforms which ranges from 0.94 to 1.0 (Table 3).

**Table 3.** The performance of Control-FREEC on NA12878 data before and after being polished by DECoNT.

| Platform | # of Events | % of Improved Events | % of Deteriorated Events | Mean Absolute Error (MAE) Difference Decreased by |
| --- | --- | --- | --- | --- |
| NovaSeq 6000 | 329 | **73.85%** | 7.59% | **0.9392** |
| HiSeq 4000 | 437 | **70.94%** | 6.86% | **1.0022** |
| BGISEQ-500 | 367 | **64.57%** | 8.17% | **0.9809** |
| MGISEQ-2000 | 367 | **64.57%** | 8.17% | **0.9809** |

We evaluate caller performance on NA12878 data obtained using four different sequencing platforms: (i) NovaSeq 6000; (ii) HiSeq 4000; (iii) BGISEQ-500; (iv) MGISEQ-2000. Note that DECoNT models did not train on the data sequenced with any of these sequencing platforms or on NA12878 sequencing data of any form. Table shows the number of CNVs reported on each sample, the percentage of improved and deteriorated events, the average decrease in absolute error after being polished by DECoNT. In all comparisons, DECoNT provides substantial improvements showing the generalizability of our models trained on 1000 Genomes data set. 1000 Genomes data set WGS samples are used as ground truth calls.

We note that the improvements provided by DECoNT on the BGI and MGI platforms are important as these systems belong to a completely different manufacturer. Since these platforms are expected to have different systematic biases in read depth distributions compared to the training data of DECoNT, we would also expect a lower testing performance. Yet, DECoNT is able to generalize well and consistently proves to be useful across a diverse set of technologies. Overall, the performance is on par with the tests obtained on Illumina Genome Analyzer II and Illumina HiSeq 2000. Polishing procedure consistently improves the performance in a platform-independent manner.

### 2.4 Polishing Performance on Calls From Unseen CNV Callers

A distinct DECoNT model is trained for every WES-based germline CNV caller. This makes sense as the call regions and numbers substantially differ among algorithms in their recommended settings (e.g., CODEX2 calls 10× more events than XHMM). We check if a DECoNT model trained using calls made by one algorithm can be used to polish the calls made by others in the absence of a trained model (e.g., due to time constraints in training).

We use the same DECoNT models trained for XHMM, CoNIFER, and CODEX2 on 1000 Genomes Project data. For each tool-specific DECoNT model, we polish the calls made by others on samples not seen during training. For instance, we polish the calls made by CODEX2, using the DECoNT model trained on XHMM calls. This experiment results in 6 tests (i.e., for two-way comparison among every tool pair). We measure the performance of the polishing procedure using duplication call precision, deletion call precision and overall precision to obtain 18 performance results in total (Methods).

We observe that DECoNT improves the performance metric in 10 out of the 18 comparisons, XHMM-trained DECoNT consistently improves the other tools’ performance in all metrics, except DEL precision when polishing calls reported by CoNIFER - ranging from 2% to 13% (Figure 3 and Supplementary Figure 1). Duplication precision is improved in most of the cases with the exception of CoNIFER-trained and CODEX2-trained DECoNT models deteriorating the performance of XHMM by 11% and 8% respectively. For deletion precision, this is not the case. Deletion precision is improved for CODEX2 for both DECoNT models. However, for CoNIFER, deletion precision is deteriorated by 13% and 45% when polished with XHMM-trained and CODEX2-trained DECoNT models respectively. This is due to very limited number of deletion CNV predictions of CoNIFER as even a small perturbation to the true positives of deletion calls yield large differences in precision. Also, CoNIFER-trained DECoNT model very slightly deteriorates deletion precision of XHMM calls by 5%. While XHMM improves overall precision for both other methods, in half of the overall precision comparisons, the performance is decreased.

**Fig. 3.**
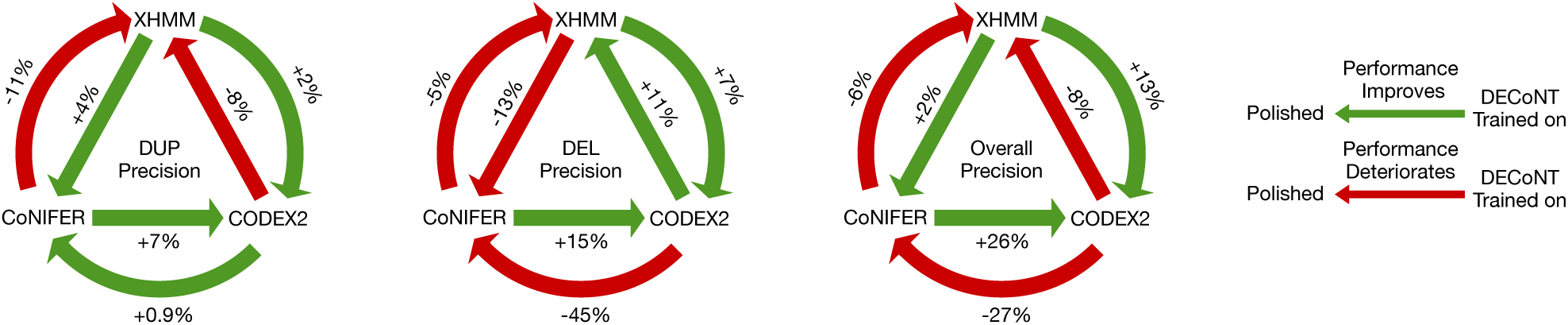
Performance of DECoNT when polishing calls from unseen CNV callers. DECoNT learns a different set of weights and a different model for each WES based CNV caller. In order to demonstrate the cross-model performance, we used DECoNT to correct CNV calls made by tools other than the ones used for training. We try every pair combination. Tools being pointed by an arrow are call generating tools (i.e., being corrected). Tools at source of the arrow are the tools that are used to train the DECoNT model. Green arrows indicate improvement and red arrows indicate deterioration in the corresponding performance metric.

Overall, DECoNT is still somewhat effective despite being trained using a different call set. The training process uses the read depth information for the event regions which enables DECoNT to generalize to polish other tools. While, arguably, it can be used to polish calls generated by other tools, a DECoNT model trained on the calls of the to-be-polished WES-based caller is suggested, as the improvements in the discussed performance metrics are larger in this case, which is expected.

### 2.5 Minimizing the false calls made by DECoNT for clinical use

Despite substantial improvements over the precision of the base callers, one could argue that for downstream usage in clinic one needs to further limit the false calls. That is, only focusing on very confident calls. We investigate the affect of this on the precision of (i) raw XHMM calls and (ii) DECoNT-corrected XHMM calls. For this, we gradually adjust both XHMM and DECoNT to be more conservative in making calls to limit false positives. That is, we set a confidence threshold and each algorithm makes a call only if an event’s probability (e.g., DUP probability) exceeds this threshold. For instance, DECoNT in the default mode would call a DEL, if DEL probability is 0.34 where DUP and NO-CALL probabilities are both 0.33. Now, if we set the confidence threshold as 0.5, a call is made only if the probability of the event is *>* 0.5.

As seen in Figure 4, making XHMM more conservative unsurprisingly increases its precision. However, we observe that this cannot replace the polishing procedure. In all categories, DECoNT correction consistently dominates the precision performance of raw XHMM calls by a large margin (by ∼20%). Second, we see that it is possible to achieve *>*∼95% precision by keeping only very high confidence calls (i.e., confidence threshold ∼= 1). We also see that such high precision is not obtained by totally sacrificing recall (i.e., making only a very small number of calls) as ∼ 1, 000 calls are still kept in the most conservative setting out of 6, 832 (see the last panel in Figure 4).

**Fig. 4.**
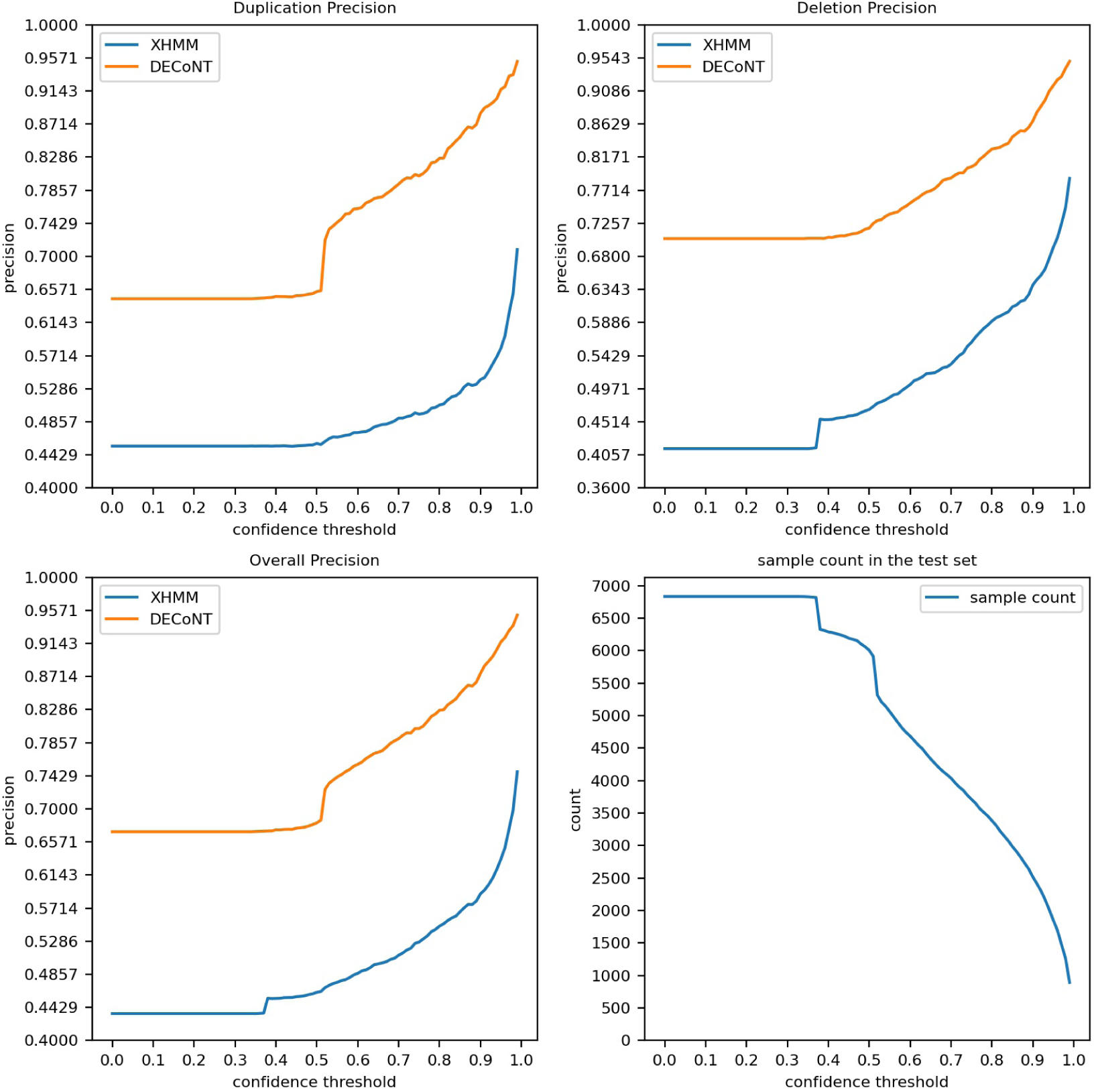
Precision of raw XHMM and DECoNT-corrected XHMM calls are shown with respect to varied confidence threshold (DEL, DUP and overall precision). 6,832 CNV calls made by XHMM on our test dataset (the 1000 Genomes WES data set test samples) are used. The ground truth is the CNV calls made by CNVnator on the corresponding WGS samples. The last panel shows the number of calls remaining as the threshold gets more conservative.

### 2.6 Genomic insights obtained from the polishing procedure

Here, we investigate the decisions made by DECoNT on the 6,832 calls made by XHMM on the 1000 Genomes test set to obtain genomic insights. First, we evaluated DECoNT for calls made on the X chromosome and pseudo-autosomal regions. We observe that the performance of XHMM is the lowest in these regions. Yet, DECoNT is still able to improve the performance in 8 out of 9 categories as shown in Supplementary Table 1.

We also analyzed the length distribution of the corrected calls. In Supplementary Figure 4, we show that true XHMM calls have a large variance in size (i.e., up to 1 Kbp). Yet, the DECoNT-corrected true calls range up to 500 kbps, showing that XHMM shows lower accuracy in shorter calls and needs polishing.

We investigated whether the importance of the DECoNT polishing varies across chromosomes. Figure 5 shows the chromosome-wise stratification of the calls where each dot represents a call made by XHMM, colored by one of the four possibilities: (i) DECoNT-polished call is correct (i.e., matching WGS call, semi ground-truth) and XHMM call is incorrect; (ii) XHMM call is correct and DECoNT agrees, (iii) Both polised and original calls are incorrect; and (iv) XHMM call is correct, DECoNT-polished call is incorrect. We observe that 80% of the XHMM calls made on chromosome 8 are changed and corrected by DECoNT. This number is 84% for chromosome Y. This indicates that researchers focusing on these regions of the genome should definitely use polishing. 44% of the XHMM calls on chromosome 20 are corrected and this is the best case scenario for XHMM, yet nearly half of the calls needed correction. We see that there is a large variance in the length of the calls made on the Y chromosome and polishing helped regardless of the length (i.e., both short and long calls are corrected). On chromosome 13, most calls are close to the median length with low variance and DECoNT almost all corrected calls are relatively short. This indicates that DECoNT is useful when the calls on a chromosome have both low and high length variance, and it can correct both short and long calls. We observe that DECoNT makes a small number mistakes and these are uniformly distributed across chromosomes.

**Fig. 5.**
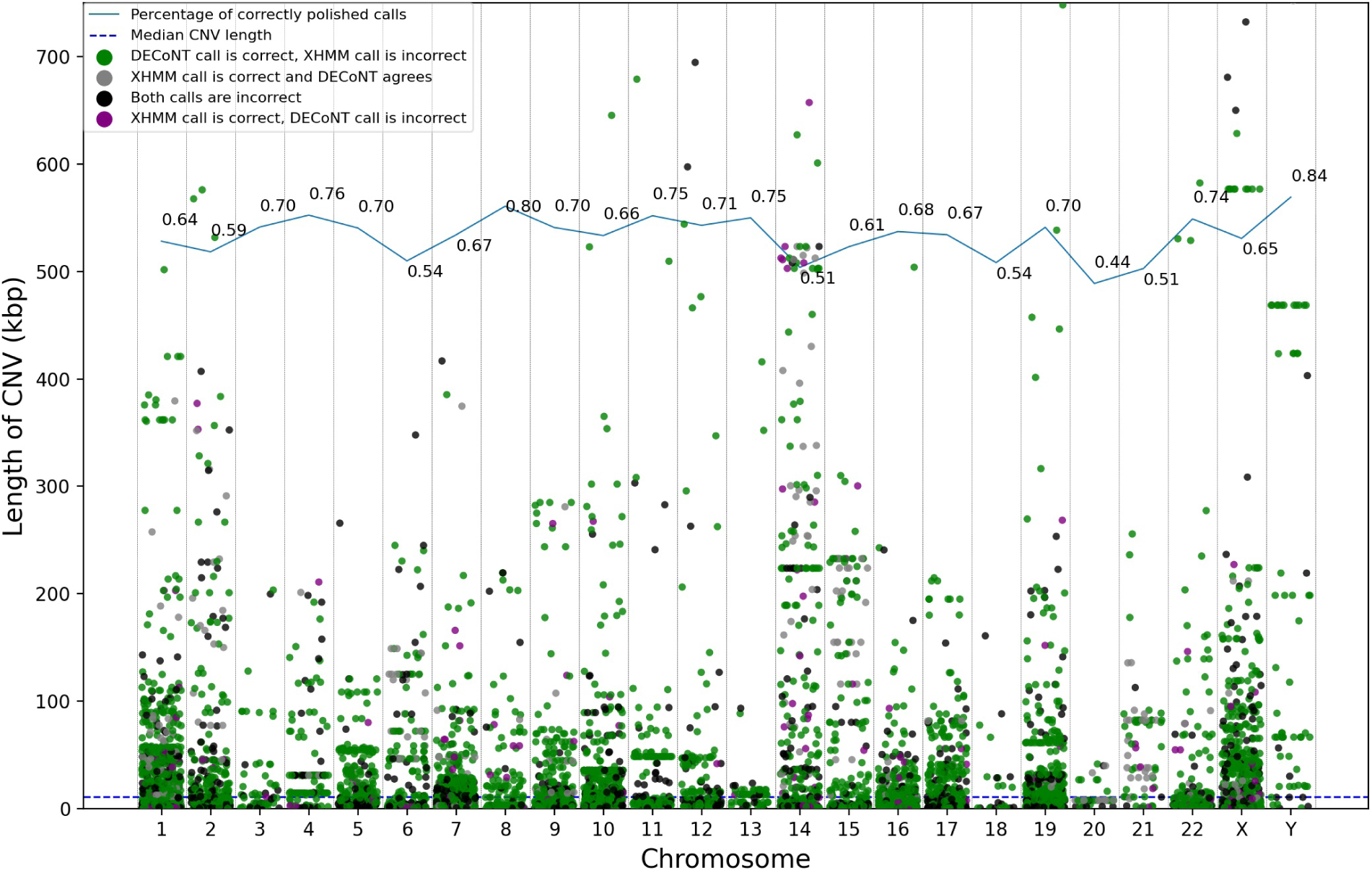
Each dot in this figure corresponds to one of the 6,832 CNV calls made by XHMM on our test dataset (the 1000 Genomes WES data set test samples). The ground truth is the CNV calls made by CNVnator on the corresponding WGS samples. The calls are stratified w.r.t. their chromosomes. The y-axis of the plot represents the length of calls. Gray dot indicates that the XHMM call is not changed by DECoNT and the prediction matches the ground truth (correct), whereas black dot indicates that both DECoNT’s and XHMM’s decisions do not match the ground truth (incorrect). Green dot indicates the call is corrected by DECoNT and XHMM call was incorrect. Finally, purple dot indicates that DECoNT changes the prediction of XHMM and both the original and the changed calls are incorrect. For each chromosome, a random jitter is added to the x-axis for better visualization. Solid line on the top of the figure shows the ratio of the number of DECoNT-corrected calls and the number of all XHMM calls in that chromosome. Dashed line indicates the median CNV call length across chromosomes.

Next, we investigated whether the calls corrected by DECoNT differ with respect to the GC content of the region. The Supplementary Figure 5 shows the GC distribution of the call regions, stratified with respect to the four categories in Figure 5. DECoNT-polished calls tend to have a larger GC content. Correctly polished calls have a mean GC% of ∼53% and correct-and-agreed XHMM calls have a mean GC% of ∼48%. XHMM has no correct calls in both GC-rich (*>*65%) and GC-poor areas (*<*35%). DECoNT is able to correct many calls in these biased regions.

Finally, we check whether the number of probes in a CNV region introduces a bias. As expected, high probe count leads XHMM to make more erroneous calls. As shown in Supplementary Figure 6, all calls with probe count larger than ∼50 were incorrect. DECoNT was able to correct all XHMM calls with probe count higher than 60. We conclude that DECoNT-polishing is essential especially in regions with systematic biases.

## 3 Discussion

High throughput sequencing platforms, since their inception in 2007, have now become the dominant source of data generation for biological and medical research and on their way to be routinely used for diagnosis and treatment guidance. Although whole human genome sequencing cost is now reduced below the $1,000 mark, whole exome sequencing will likely remain the main workhorse in clinical settings due to i) lower cost, ii) capturing almost all actionable genetic defects within exons, and iii) smaller data size that reduces computational burden for analysis. However, the main drawback of WES has been discovery and genotyping of CNVs. First, depth of coverage among exons are not uniform, making it very difficult to apply read depth based methods. Second, the reads often do not span CNV breakpoints which is a must for read pair and split read based approaches. Therefore it is often necessary to complement WES studies with alternative approaches such as array comparative genomic hybridization or quantitative RT-PCR.

We specifically designed our new algorithm, DECoNT, to address this limitation as a CNV call *polisher*. Using a deep learning approach, we were able to boost both the precision of several widely-used state-of-the-art algorithms that use WES data for CNV discovery. Although we trained DECoNT using matched WGS and WES samples from the 1000 Genomes Project, we also demonstrated that the performance gain is independent from the training data, the capture kit, and the sequencing platform. Therefore, the trained model is portable, and it can be applied to data sets regardless of the data generation protocol without requiring new samples to train DECoNT.

Copy number variation is an important cause of genetic diseases that may be difficult to characterize in clinical settings without specific assays. WES is a powerful method to genotype small mutations but so far it has been unsuccessful to discover large CNVs that have a more direct effect in gene losses. DECoNT aims to help ameliorate high false discovery rate problems related to CNV characterization using WES, also including integer copy number prediction. Therefore DECoNT adds an important type of genomic variation discovery to the capabilities of WES and enhances the genome analysis arsenal in the clinic.

DECoNT uses WGS-derived CNV calls as labels for training. Note that these labels cannot serve as the ground truth but rather as the semi-ground truth. Unfortunately, there does not exist sufficiently large hand-curated labeled data for training a model. Chaison et al. provides hand-curated CNV calls on 9 samples which let us only perform validation. While a larger sample set was used, the latest release by HGSVC [5] contains only 674 CNVs, which is a very small number for training DECoNT. This call set can also be regarded as a consensus call set (i.e., result of a consensus caller) because it is generated by using three different calling pipelines, and supported by long-read sequencing, StrandSeq, and optical mapping analysis of a subset of these genomes. We attempted training a model with this data set but unsurprisingly the model did not converge. For comparison, for training the DECoNT model for XHMM, we used ∼ 68*k* calls. While these hand curated high quality data sets and consensus callers are certainly going to help DECoNT to achieve higher precision, they are quite limited in size which currently prohibits training. We foresee that with increasing size of call sets, DECoNT’s performance will also increase. Yet, it is evident that CNV calling on WGS data is more accurate compared to CNV calling on WES data even when using a single over-the-shelf WGS-based CNV caller. Thus, DECoNT transfers these higher confidence labels into the WES domain and is limited by the precision of the underlying WGS-based CNV caller (CNVnator in this case) [1]. Abyzov et al. report that CNVnator has high sensitivity (86%-96%), and low false-discovery rate (3%-20%). This corresponds to a precision range of 80% to 97% (mean 11.8%). Note that [1] obtains these performance results on two high coverage (20× - 32×) trios in the 1000 Genomes Pilot Project data set [38]. DECoNT uses the newer 1000 Genomes data set at 30× coverage generated using NovaSeq [5]. Thus, we expect the error tolerance of CNVnator to be better or similar in our analyses.

One issue with the polisher is to come up with a recipe to set the parameters of the base caller to achieve the best performance. In this study, we mostly used the suggested parameters and had to relax CoNIFER’s parameters as it hardly returned any calls. One other option is to run the base caller in the most liberal setting to improve sensitivity and let DECoNT correct the likely higher number of false positives. We tested if this is feasible using XHMM which is the best performing algorithm in our benchmarks. As detailed in Supplementary Note 3, we ran it also in a liberal and a conservative setting and then polished the calls using DECoNT. The precision improvement of tool was stable at ∼ 22% in all three settings. The polished precision of the liberal setting was worse by 10% compared to the suggested setting. Conservative and suggested setting precision values differed by only ∼ 1%. Thus, we suggest using the *suggested* parameter settings of the base caller unless it hardly makes calls and provides an insufficient number of calls for training.

The next challenge will be relieving DECoNT from dependence on existing variation callers, and make it a standalone, highly accurate CNV discovery tool using whole exome sequencing. One possible such direction for DECoNT could be redesigning the architecture to work with bin-level data. That is, most baseline callers first analyze read depth in small bins and then, adjacent bins are smoothed/combined via a segmentation algorithm which are returned as final calls if enough evidence exists along multiple neighboring bins. Currently, DECoNT-the-polisher, is working with this final decision which is limiting. Working with bin level data will require an architecture that can handle one basepair resolution whereas now DECoNT works with kilobase-sized windows which are averaged. As is, the Bi-LSTM component is prone to vanishing/exploding gradients due to possibly very long read depth sequences.

## 4 Methods

### 4.1 Data set

For training and testing of DECoNT, we used 1, 000 samples (i.e., HG00096 to HG02356, when sample IDs are alphabetically ordered) from the 1000 Genomes Project [5]. For these samples we obtain both WES and WGS data. WES samples were captured using the NimbleGen SeqCap EZ Exome v3 as capture kit, and sequenced to an average of 50× depth with Illumina Genome Analyzer II and Illumina HiSeq 2000 platforms. The average read length is 76 bps. Reads were aligned to the GRCh38 using the BWA-MEM aligner [25]. WGS samples were also sequenced using the same platforms with an average read length of 100bps. Average depth coverage for this set is 30×. For XHMM, CoNIFER and CODEX2, the ground truth CNV calls are obtained using CNVnator [1] tool. For Control-FREEC and CNVkit, the ground truth exact copy number variation events are obtained using mrCaNaVaR [2].

For tools that output a categorical prediction of a CNV, we also use a highly validated CNV call set published in Chaisson *et al*., [6] as another validation source. The WGS CNV calls in this call set are thoroughly validated. That is, they were obtained via a consensus of 15 different WGS CNV callers with comparisons against high quality PB-SVs that have single base breakpoint resolution. We obtain WGS CNV calls for these 9 samples from 1000 Genomes data set. (i.e., HG00512, HG00513, HG00514, HG00731, HG00732, HG00733, NA19238, NA19239, NA19240). We also obtained aligned WES reads of these samples, with the exception of HG00514 for which no WES data was available. This data set is only used for testing.

### 4.2 DECoNT Model

#### Problem Formulation

Let *X* denote the set of CNV events detected on the WES data set by a WES-based CNV caller, and *X*^(*i*)^ denote the *i*^*th*^ event. *F*_*i*_ denotes the set of features we use for *X*^(*i*)^ which contains the following information: (i) the chromosome that the CNV event occured 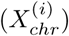; (ii) the start coordinate of the CNV event 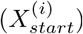 ; (iii) the end coordinate of the CNV event; (iv) the type (e.g., deletion) of the called event 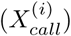; and (v) the read depth vector between 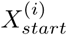 to 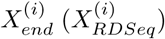. Let 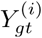 denote the ground truth label obtained from the WGS CNV call for *X*^(*i*)^. There are two cases: (i) For the tools that predict the existence of an event 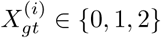, denoting no call, deletion or duplication, respectively; and (ii) For the tools that predict the copy number 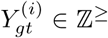. Then, the problem at hand is formulated as a classification task for (i), and as a regression task for (ii). That is, our goal is to learn a function *f* such that 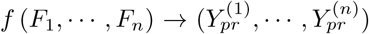 such that the difference is between (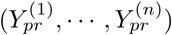) and (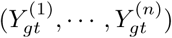) is minimized with respect to a loss function. Here, *n* = |*X*| and 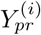 is the predicted label for *X*^(*i*)^ and it is in the same domain as 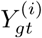 in respective tasks.

#### DECoNT Architecture

DECoNT is an end-to-end multi-input neural network designed for polishing and improving the performance of the WES-based germline CNV callers. It is capable of improving accuracy of WES CNV calling for both exact CNV prediction (i.e., integer) and categorical CNV prediction cases (i.e., deletion, duplication or no call). For each CNV caller, a distinct network is trained.

DECoNT’s pipeline for the categorical CNV prediction case can be divided into three main building blocks: (i) a data preprocessing step that extracts the read depth for genomic regions of interest (i.e., CNV call regions made by the CNV caller). It also normalizes the read depth sequence and acts as a regularizer for the model. Resulting read depth information is −1 padded to the length of the longest call sequence and masked; (ii) bidirectional LSTM network (BiLSTM) that inputs the read depth sequence and extracts the required encoded features (i.e., embeddings). This subnetwork has 128 neurons in each direction and is followed by a batch normalization layer; and (iii) a 2-layered fully connected (FC) neural network that inputs the embedding calculated by Bi-LSTM, concatenated with the prior CNV prediction of the CNV caller (a one-hot-encoded vector). The first FC layer has 100 neurons and uses ReLU activation. The output layer has 3 neurons and it calculates the posterior probability of each event via softmax activation: no call, deletion, or duplication. We use weighted cross-entropy as the loss function. This architecture has a total of 160, 351 parameters with 159, 837 of which are trainable. The rest are the batch normalization parameters.

For a training data set of *N* samples, the formulation of DECoNT can be summarized as follows:

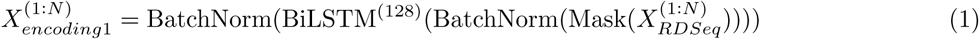

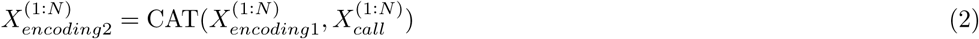

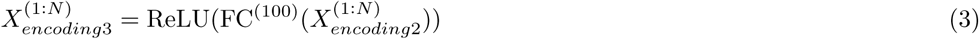

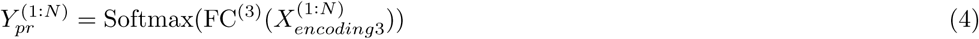

where BiLSTM^(·)^ represents bi-direcitonal LSTM layer with · hidden units in each direction. Similarly, FC^(·)^ represents a dense layer with · neurons. ReLU and BatchNorm stand for rectified linear unit activation function and batch normalization respectively.

Using 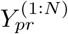 and 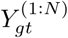 training phase minimizes the categorical cross-entropy loss. We use Adam optimizer [20] with a mini batch size of 128 samples. All weights in the network are initialized using Xavier initialization [10].

DECoNT’s pipeline for the exact (i.e., integer) CNV prediction is almost the same as the one described above. The first difference is instead of taking the one-hot encoded version of the CNV call, it inputs an integer value representing the called copy number. The second difference is at the output layer. Instead of 3 neurons with softmax activation, this version has a single neuron with ReLU activation to perform regression instead of classification. It has a total of 160, 149 parameters, 159, 635 of which are trainable. Again, the rest are the batch normalization parameters. So, the last layer in the formulation above (Eq. 4) is replaced by the following layer and in this case 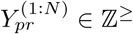.

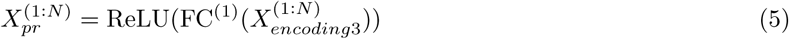

Using 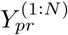 and 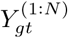 training phase now minimizes the mean absolute error loss. Again, we use Adam optimizer with a mini batch size of 128 samples and we use Xavier initialization for weights.

### 4.3 Polishing the State-of-the-art WES-based Germline CNV Callers

We polish the CNV calls made by four state-of-the-art WES-based germline CNV callers (i) XHMM [9], (ii) CoNIFER [21], (iii) CODEX2 [16], (iv) Control-FREEC [4], and (v) CNVkit [35]. Tools (i - iii) perform categorical CNV prediction and (iv) and (v) perform exact CNV prediction. We use calls made on the WGS samples by CNVnator [1] as the ground truth call set for discrete predictions and the exact copy number predictions made by mrCaNaVaR as the ground truth call set for integral prediction (i.e., Control-FREEC). First, DECoNT obtains the results of these tools (i - iv). Then, it learns to correct these calls on a portion of the 1000 Genomes data set using ground truth calls. Finally, on the left out test portion of the data, we compare the performance of the CNV callers before and after polishing by DECoNT.

#### Settings for the WES-based CNV Callers

We follow the recommended settings for the WES-based Callers. For XHMM, the parameters are set as follows: (i) *Pr*(start DEL) = *Pr*(start DUP) = 1*e* − 08, (ii) mean number of targets in CNV (geometric distribution) = 6, (iii) mean distance between targets within CNV (exponential decay) = 70*kb*, and (iv) DEL, DIP, DUP read depth distributions modeled as ∼𝒩(−3, 1), ∼𝒩(0, 1) and ∼𝒩(3, 1), respectively. Also, for XHMM nBins parameter is set as 200 which is the default setting. For CODEX2, minimum read coverage of 20 was enforced at the filtering step. Then, the algorithm automatically chooses its parameter, *K*, using BIC (i.e. Bayesian Information Criterion) and AIC (i.e. Akaike Information Criterion). CoNIFER performs SVD on the data matrix and then removes *n* singular vectors with *n* largest singular values. We set *n* to 6. Control-FREEC has 45 parameters which were all set to default values as stated in [4]. CNVkit uses a rolling median technique to recenter each on- or off-target bin with other bins of similar GC content, repetitiveness, target size or distance from other targets, independently of genomic location [35]. We used recommended settings for CNVkit as well where log2 read depth below threshold = −5, above threshold = 1.0.

#### Training Settings for DECoNT

We train a DECoNT model for each of the above-mentioned tools. The set *X* of CNV calls per tool is shuffled and divided into training, validation and testing sets which contain 70%, 20%, and 10% of the data, respectively. The number of events in the test sets are 6, 832 (3101 no-calls, 2098 duplications, 1633 deletions); 81, 761 (67885 no-calls, 3042 duplications, 10834 deletions); 180 (85 no-calls, 43 duplications, 52 deletions); 20, 482 (minimum copy number is 0, maximum copy number is 585); 39720 (minimum copy number is 34, maximum copy number is 0) for XHMM, CODEX2, CoNIFER, Control-FREEC, and CNVkit, respectively. The second input of the algorithm is the read depth for the CNV-associated regions on the WES data. We calculate it using the Sambamba tool [37]. For all tools other than CODEX2, DECoNT is trained up to 30 epochs with early stopping by checking the loss on the validation fold. Training for CODEX2 has a maximum epoch number 60. For training, DECoNT uses final CNV calls (i.e., concatenated bin-level calls) made by the CNV callers.

#### Performance Metrics

Tools (i - iii) predict CNVs either as deletion or duplication. The main problem of these callers are false discovery rates [42,36]. Given a deletion or duplication call by tools (i - iii), DECoNT outputs a probability for the call to be deletion, duplication or no call (i.e., false discovery). The option with the highest probability is returned as the prediction.

In order to assess the performance of tools (i - iii) before and after being polished, we calculate the following performance metrics using 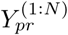 and 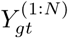 : (i) duplication call precision; (ii) deletion call precision, and (iii) overall precision. We first define the following variables: TP_1_ := number of duplications correctly identified; TP_2_ := number of deletions correctly identified; FP_1_ := number of duplications incorrectly identified; FP_2_ := number of deletions incorrectly identified.

Then, the performance metrics are defined as follows:

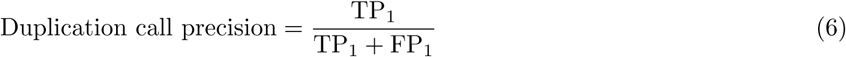

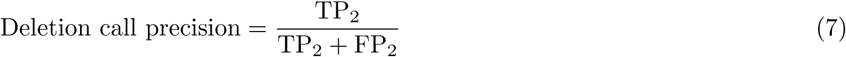

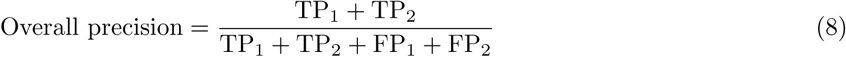

In order to test DECoNT’s performance on exact CNV prediction problem, which is a regression task, we use Absolute Error (*AE*) between the predicted and ground truth copy number values. For an event *X*_*i*_, *AE*^(*i*)^ is defined as follows:

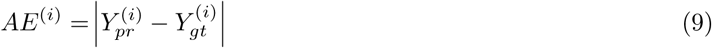

#### Time Performance

All models are trained on a SuperMicro SuperServer 4029GP-TRT with 2 Intel Xeon Gold 6140 Processors (2.3GHz, 24.75M cache), 251GB RAM, 3 NVIDIA GeForce RTX 2080 Ti (11GB, 352Bit) and 1 NVIDIA TITAN RTX GPUs (24GB, 384Bit). We used 4 GPUs in parallel to train all 5 models and total training times were approximately as follows: ∼ 70, 12, 95, 50, and 20 hours for XHMM, CoNIFER, CODEX2, Control-FREEC, and CNVkit, respectively. Note that training is performed offline. The average polishing time per sample is in the order of seconds, for all models.

### 4.4 Polishing Samples from Other Sequencing Platforms

The training data we use is obtained using Illumina Genome Analyzer II and Illumina HiSeq 2000 machines. We check if models trained on these 1000 Genomes data can be used to polish CNV calls made on WES samples obtained using other sequencing platforms or capture kits that have not been seen by DECoNT.

We obtain the WES data for the sample NA12878, sequenced using four different platforms: (i) Illumina NovaSeq 6000; (ii) Illumina HiSeq 4000; (iii) BGISEQ-500; and (iv) MGISEQ-2000. Reads are aligned to the reference genome (GRCh38) using BWA [25] with *-mem* option and default parameters. Average depth coverage for these samples are 241×, 395×, 328×, and 129×, respectively. We use these four samples only for testing. All considered WES-based CNV callers are used to call CNV events on these four WES samples with default parameters. Using the CNVnator calls obtained on the WGS sample for NA12878 as the ground truth, we measure the performance the CNV callers before and after polishing with DECoNT. Note that NA12878 data is not included in the training data set in any form.

### 4.5 Polishing Other WES-based CNV Caller Algorithms

In our framework, a separate DECoNT model is trained for every WES-based germline CNV caller. We check if a DECoNT model trained using calls made by one algorithm can be used to polish the calls made by others in the absence of a trained model.

We use the same three models trained with the settings described in Section 4.3 for XHMM, CoNIFER, and CODEX2. For each tool-specific DECoNT model, we polish the calls made by others. Here the training and testing folds are again exclusive. For testing, we use the same test folds for each tool as described in in Section 4.3 This experiment results in 6 tests (i.e., for two-way comparison among every tool pair). We measure the performance of the polishing procedure using duplication precision, deletion precision and overall precision to obtain 18 performance results in total.

## Author Contributions

AEC and CA designed and supervised the study. AEC and FO designed the model. FO implemented the software and performed the experiments. AEC, CA and FO wrote the manuscript.

## Data Availability

The training data used for DECoNT are obtained from 1000 Genomes Project. The WES bam files are available at ftp://ftp.1000genomes.ebi.ac.uk/vol1/ftp/data_collections/1000_genomes_project/data/. WGS CNV calls are obtained from CNVnator tool, they are already processed and made available at: ftp://ftp.1000genomes.ebi.ac.uk/vol1/ftp/data_collections/1000G_2504_high_coverage_SV/working/20190825_Yale_CNVnator/. WES reads for NA12878 sample were obtained from: https://www.ncbi.nlm.nih.gov/sra with accession codes SRX5191370, SRX5191369, SRX5180030, and SRX5180221 for NovaSeq 6000, HiSeq 4000, BGISEQ-500, and MGISEQ-2000, respectively. Highly validated WGS CNV calls presented in Chaisson et. al. obtained from dbVar https://www.ncbi.nlm.nih.gov/dbvar/ with accession nstd152. The inputs we use (i) CNV calls made by third party CNV callers and (ii) the calculated read depth data are also available at https://zenodo.org/record/3865380 inside respective folders of the analysis. All other data that support the key findings of this paper can be found in the article and corresponding supplementary tables as referenced in the text.

## Code Availability

DECoNT is implemented and released at https://github.com/ciceklab/DECoNT under CC-BY-NC-ND 4.0 International license. All custom python scripts that were used to generate matched WGS and WES CNV data are also available on the GitHub page. The scripts used to generate the data for all figures and tables in the manuscript are provided at https://zenodo.org/record/3865380.

## Acknowledgments

We would like to thank Vijay Kumar and Santosh Girirajan for their help with obtaining results on CNLearn. The following cell lines/DNA samples were obtained from the NIGMS Human Genetic Cell Repository at the Coriell Institute for Medical Research: [NA06984, NA06985, NA06986, NA06989, NA06994, NA07000, NA07037, NA07048, NA07051, NA07056, NA07347, NA07357, NA10847, NA10851, NA11829, NA11830, NA11831, NA11832, NA11840, NA11843, NA11881, NA11892, NA11893, NA11894, NA11918, NA11919, NA11920, NA11930. NA11931, NA11932, NA11933, NA11992, NA11994, NA11995, NA12003, NA12004, NA12005, NA12006, NA12043, NA12044, NA12045, NA12046, NA12058, NA12144, NA12154, NA12155, NA12156, NA12234, NA12249, NA12272, NA12273, NA12275, NA12282, NA12283, NA12286, NA12287, NA12340, NA12341, NA12342, NA12347, NA12348, NA12383, NA12399, NA12400, NA12413,, NA12414, NA12489, NA12546, NA12716, NA12717, NA12718, NA12748, NA12749, NA12750, NA12751, NA12760, NA12761, NA12762, NA12763, NA12775, NA12776, NA12777, NA12778, NA12812, NA12813, NA12814, NA12815, NA12827, NA12828, NA12829, NA12830, NA12842, NA12843, NA12872, NA12873, NA12874, NA12878, NA12889, NA12890]. These data were generated at the New York Genome Center with funds provided by NHGRI Grant 3UM1HG008901-03S1. The corresponding data is released by the HGSVC [5].

