## Supplementary Material for "Polishing Copy Number Variant Calls on Exome Sequencing Data via Deep Learning"

### Supplementary Material
### for
### Polishing Copy Number Variant Calls on Exome Sequencing Data via Deep Learning

#### 1 Supplementary Figures

| | | Unpolished | | | DECoNT-XHMM Polished | | | DECoNT-CoNIFER Polished | | | DECoNT-CODEX2 Polished | | |
|---|---|---|---|---|---|---|---|---|---|---|---|---|---|
| | | NO CALL | DUP | DEL | NO CALL | DUP | DEL | NO CALL | DUP | DEL | NO CALL | DEL | DEL |
| XHMM Ground Truth | NO CALL | NA | 1708 | 1447 | 1974 | 790 | 391 | 2104 | 310 | 741 | 2647 | 505 | 3 |
| | DUP | NA | 1587 | 508 | 360 | 1589 | 146 | 1384 | 263 | 448 | 1666 | 422 | 7 |
| | DEL | NA | 198 | 1384 | 217 | 84 | 1281 | 715 | 208 | 659 | 1359 | 212 | 11 |
| CoNIFER Ground Truth | NO CALL | NA | 77 | 8 | 15 | 53 | 17 | 76 | 3 | 6 | 69 | 15 | 1 |
| | DUP | NA | 39 | 4 | 2 | 31 | 4 | 5 | 27 | 11 | 30 | 12 | 1 |
| | DEL | NA | 42 | 10 | 4 | 24 | 10 | 9 | 6 | 37 | 32 | 20 | 0 |
| CODEX2 Ground Truth | NO CALL | NA | 7955 | 6413 | 5337 | 3028 | 6003 | 1248 | 166 | 958 | 10974 | 1773 | 1621 |
| | DUP | NA | 2081 | 1786 | 368 | 1145 | 2354 | 339 | 130 | 561 | 896 | 1794 | 1177 |
| | DEL | NA | 7240 | 6774 | 1015 | 3723 | 9276 | 945 | 371 | 2282 | 3662 | 4216 | 6136 |

**Supplementary Figure 1.** The confusion matrices of the WES-based CNV callers before and after polishing with DECoNT on 1000 Genomes Data test samples. Confusion matrices given with blue borders represent unpolished predictions of corresponding WES-based CNV tools. Since DECoNT only operates on the calls made by a CNV caller, the first column for each unpolished confusion matrix is set as NA (i.e. Not Applicable). The red-bordered confusion matrices are the polished versions of a CNV caller with a DECoNT model trained on the calls made by the same caller (to produce Fig 2). Other confusion matrices are polised version of the CNV caller corrected by a DECoNT model trained on the calls made by a different caller (to produce Fig. 3). Notice the decrease in the number of false positives for both deletion and duplication calls in all platforms.

| | | Unpolished | | | Polished | | |
|---|---|---|---|---|---|---|---|
| | | NO CALL | DUP | DEL | NO CALL | DUP | DEL |
| XHMM Ground Truth | NO CALL | NA | 352 | 210 | 195 | 264 | 103 |
| | DUP | NA | 34 | 18 | 18 | 28 | 6 |
| | DEL | NA | 144 | 79 | 55 | 99 | 69 |
| CoNIFER Ground Truth | NO CALL | NA | 79 | 0 | 46 | 13 | 20 |
| | DUP | NA | 11 | 0 | 3 | 4 | 4 |
| | DEL | NA | 32 | 0 | 13 | 8 | 11 |
| CODEX2 Ground Truth | NO CALL | NA | 3251 | 2472 | 4643 | 588 | 492 |
| | DUP | NA | 145 | 118 | 128 | 87 | 48 |
| | DEL | NA | 1973 | 1636 | 1225 | 1209 | 1175 |

**Supplementary Figure 2.** Confusion matrices of the WES-based CNV callers before and after polishing with DECoNT on highly validated CNV callset published in Chaisson et. al. [1]. Similar to Fig. 1 tool provides great false discovery correction with slight true positive deterioration for both deletion and duplication calls, yielding much better performance metric results.

| | | NovaSeq6000 | | | | | | HiSeq4000 | | | | | | BGI500 | | | | | | MGI2000 | | | | | |
|---|---|---|---|---|---|---|---|---|---|---|---|---|---|---|---|---|---|---|---|---|---|---|---|---|---|
| | | Unpolished | | | Polished | | | Unpolished | | | Polished | | | Unpolished | | | Polished | | | Unpolished | | | Polished | | |
| | | NO CALL | DUP | DEL | NO CALL | DUP | DEL | NO CALL | DUP | DEL | NO CALL | DUP | DEL | NO CALL | DUP | DEL | NO CALL | DUP | DEL | NO CALL | DUP | DEL | NO CALL | DUP | DEL |
| XHMM Ground Truth | NO CALL | NA | 0 | 65 | 24 | 4 | 37 | NA | 0 | 79 | 27 | 6 | 46 | NA | 21 | 10 | 10 | 12 | 9 | NA | 21 | 10 | 10 | 12 | 9 |
| | DUP | NA | 2 | 17 | 6 | 2 | 11 | NA | 1 | 18 | 10 | 1 | 8 | NA | 1 | 6 | 1 | 1 | 5 | NA | 1 | 6 | 1 | 1 | 5 |
| | DEL | NA | 1 | 7 | 2 | 0 | 6 | NA | 1 | 10 | 0 | 1 | 10 | NA | 0 | 3 | 0 | 0 | 3 | NA | 0 | 3 | 0 | 0 | 3 |
| CoNIFER Ground Truth | NO CALL | NA | 0 | 0 | 0 | 0 | 0 | NA | 0 | 24 | 9 | 1 | 14 | NA | 0 | 5299 | 3088 | 846 | 1365 | NA | 0 | 5299 | 3088 | 846 | 1365 |
| | DUP | NA | 0 | 0 | 0 | 0 | 0 | NA | 0 | 14 | 6 | 1 | 7 | NA | 0 | 67 | 23 | 10 | 34 | NA | 0 | 67 | 23 | 10 | 34 |
| | DEL | NA | 0 | 0 | 0 | 0 | 0 | NA | 0 | 9 | 4 | 0 | 5 | NA | 0 | 299 | 115 | 58 | 126 | NA | 0 | 299 | 115 | 58 | 126 |
| CODEX2 Ground Truth | NO CALL | NA | 613 | 442 | 840 | 105 | 110 | NA | 503 | 377 | 607 | 164 | 109 | NA | 320 | 250 | 478 | 33 | 59 | NA | 320 | 250 | 478 | 33 | 59 |
| | DUP | NA | 32 | 33 | 14 | 31 | 20 | NA | 19 | 19 | 9 | 19 | 10 | NA | 21 | 13 | 12 | 13 | 9 | NA | 21 | 13 | 12 | 13 | 9 |
| | DEL | NA | 98 | 118 | 43 | 87 | 86 | NA | 107 | 92 | 54 | 69 | 76 | NA | 63 | 72 | 32 | 37 | 66 | NA | 63 | 72 | 32 | 37 | 66 |

**Supplementary Figure 3.** Confusion matrices of the WES-based CNV callers before and after polishing with DECoNT on NA12878 data obtained from different sequencing platforms: (i) NovaSeq6000; (ii) HiSeq4000; (iii) BGI500; (iv) MGI2000. Since DECoNT only operates on the calls made by a CNV caller, the first column for each unpolished confusion matrix is set as NA (i.e. Not Applicable). Since CoNIFER does not report any calls on NovaSeq6000 platform, DECoNT has no input to polish and thus the comparison is not applicable. Similar to Figures 1 and 2, we observe that DECoNT substantially decreases the number of false discoveries with slight true positive deterioration for both deletion and duplication calls.

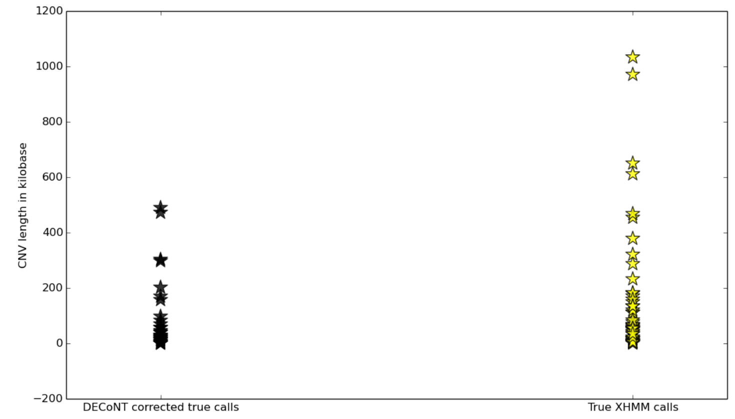


**Supplementary Figure 4.** This figure shows the length distribution of true raw XHMM calls and true DECoNT-corrected XHMM calls obtained on the 1000 Genomes WES data set test samples. The ground truth is the CNV calls made by CNVnator on the corresponding WGS samples. We see that for smaller size CNVs XHMM requires more correction by DECoNT. However, again, vast majority of the CNVs cannot be distinguished by the CNV length to decide whether it needs a DECoNT correction.

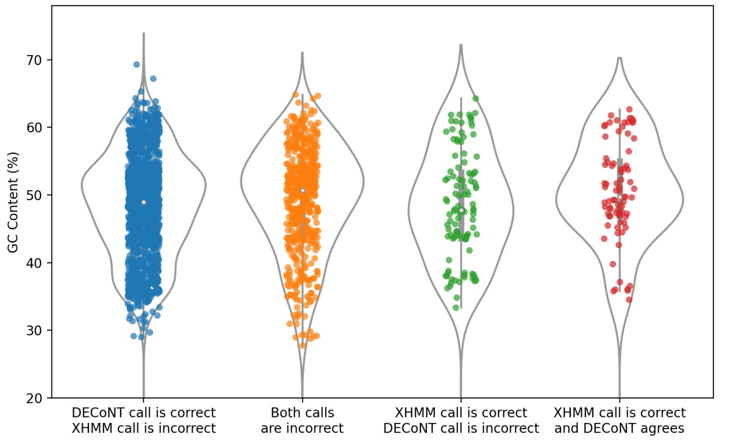


**Supplementary Figure 5.** This figure shows the probe content distribution of the XHMM calls obtained on the 1000 Genomes WES data set test samples. The ground truth is the CNV calls made by CNVnator on the corresponding WGS samples. The ground truth is the CNV calls made by CNVnator on the corresponding WGS samples. Blue dots indicate that the original XHMM call is changed by DECoNT and the changed prediction matches the ground truth (correct). Green dots indicate that the original XHMM call is changed by DECoNT and the changed prediction does not match the ground truth (incorrect). Red dots indicate original XHMM call is correct and DECoNT agreed. Finally, yellow dots indicate that both DECoNT's and XHMM's calls are incorrect. For each category, a random jitter is added to the x-axis for better visualization.

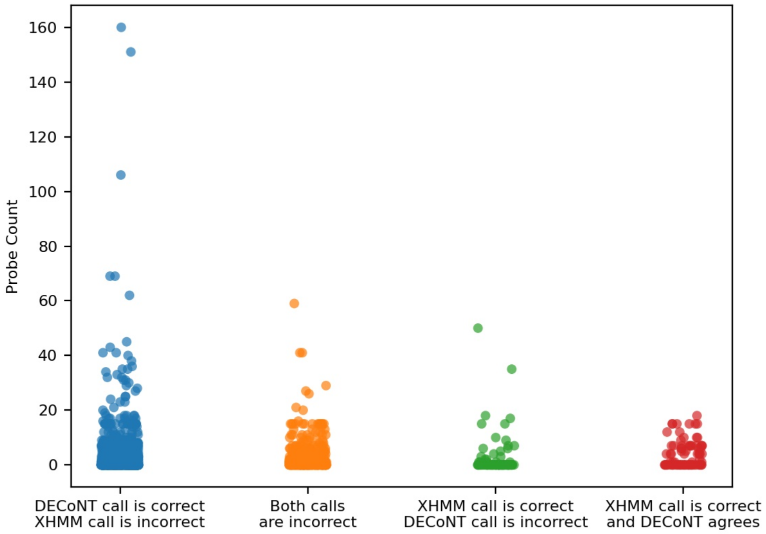


**Supplementary Figure 6.** This figure shows the probe content distribution of the XHMM calls obtained on the 1000 Genomes WES data set test samples. The ground truth is the CNV calls made by CNVnator on the corresponding WGS samples. The ground truth is the CNV calls made by CNVnator on the corresponding WGS samples. Blue dots indicate that the original XHMM call is changed by DECoNT and the changed prediction matches the ground truth (correct). Green dots indicate that the original XHMM call is changed by DECoNT and the changed prediction does not match the ground truth (incorrect). Red dots indicate original XHMM call is correct and DECoNT agreed. Finally, yellow dots indicate that both DECoNT's and XHMM's calls are incorrect. For each category, a random jitter is added to the x-axis for better visualization.

#### 2 Supplementary Tables

**Supplementary Table 1.** This table summarizes the polishing performance of DECoNT on the X chromosome, PAR1 and PAR2 regions of the males in the test split obtained from 1000 Genomes WES samples. The base caller in this analysis is XHMM. The results are obtained on the test samples from the 1000 Genomes dataset and the ground truth is obtained from the CNVnator calls on WGS of the same samples.

| | Deletion Precision (Unpolished-Polished) | Duplication Precision (Unpolished-Polished) | Overall Precision (Unpolished-Polished) |
|---|---|---|---|
| X Chromosome | 0.0350 - 0.1153 | 0.3018 - 0.4753 | 0.1702 - 0.4072 |
| PAR1 | 0.1667 - 0.2500 | 0.7083 - 0.6112 | 0.5278 - 0.5455 |
| PAR2 | 0.3334 - 0.6667 | 0.25 - 0.50 | 0.2871 - 0.4 |

**Supplementary Table 2.** In addition to the WES CNV Callers presented in the manuscript, we have also trained a DECoNT model for CNVKit on, again, 1000 Genomes WES samples, using the CNVnator calls obtained on WGS data as ground truth.

| CNVKit 1000 Genomes Ground truth call-set is 1000 Genomes WGS samples. | Deletion Precision | Duplication Precision | Overall Precision |
|---|---|---|---|
| Before DECoNT | 0.0940 | 0.1525 | 0.1234 |
| After DECoNT | 0.1234 | 0.5497 | 0.2527 |

**Supplementary Table 3.** The 8 WES CNV calls that DECoNT and CNLearn does not agree are presented. The ground truth CNV calls are obtained through CNVNator WGS CNV calls. Note that, CNLearn samples are polished with a DECoNT model trained with XHMM data. Training a DECoNT model with consensus calls made by CNLearn would increase performance.

| Sample | Chromosome | CNV Start | CNV End | CNLearn Prediction | DECoNT Prediction | Ground Truth (CNVNator WGS Calls) |
|---|---|---|---|---|---|---|
| NA19144 | 11 | 6128771 | 6170380 | DEL | NO-CALL | NO-CALL |
| NA19144 | chr14 | 73541473 | 73573608 | DUP | DEL | NO-CALL |
| NA11832 | chr6 | 32519300 | 32666612 | DUP | DEL | DEL |
| NA11832 | chr15 | 34386562 | 34528116 | DUP | DEL | DUP |
| NA18968 | chr6 | 29889285 | 29945317 | DUP | DEL | DEL |
| NA18968 | chr6 | 32519300 | 32579157 | DUP | DEL | DEL |
| NA18968 | chr6 | 32584060 | 32665112 | DUP | NO-CALL | DEL |
| NA12249 | chr16 | 55810440 | 55826342 | DUP | NO-CALL | DUP |

#### 3 Supplementary Notes

**Supplementary Note 1** For the integer CNV calls of Control-FREEC, we have categorized the calls such that Copy Number > 2 is Duplication, Copy Number < 2 is Deletion and Copy Number = 2 is No-Call. Then, we evaluated the polishing performance with the performance metrics defined in Section 4.3 Performance Metrics with respect to the 1000 Genomes WGS CNV calls of CNVNator:

- Duplication Precision was increased from 0.1063 to 0.3932
- Deletion Precision was increased from 0.2578 to 0.5936
- Overall Precision was increased from 0.1277 to 0.4432

**Supplementary Note 2** In order to show the need for a complex machine learning model like DECoNT for this polishing task, we also experimented with traditional machine learning methods such as Support Vector Machines (SVM), Logistic Regression and Polynomial Regression (degree = 2) as polishers. We used the scikit-learn implementations and the default parameters. These algorithms are run with the same settings we used for DECoNT. We worked on the 1000 Genomes dataset samples and same the train-test split. We input the same features into these models as we input to DECoNT: read depth and the call of the baseline caller. We used the corresponding WGS calls by CNVnator as the ground truth as we do for DECoNT. We used XHMM and FREEC as the baseline callers for this experiment.

Below, we show that these models actually cannot polish the calls and deteriorate the results. See the notes below:

- XHMM predictions result in 0.4541 and 0.4144 precision for duplication and deletion calls, respectively.
- When correcting XHMM calls, SVM based model predictions result in 0.3562 and 0.3321 in precision for duplication and deletion calls, respectively.
- When correcting XHMM calls, Logistic Regression based model predictions result in 0.3334 and 0.2174 in precision for duplication and deletion calls, respectively.

- Control-FREEC predictions result in a MSE of 37.17 with standard deviation of 75.89
- When correcting Control-FREEC calls, Polynomial Regression polished model predictions result in a MSE of 58.10 with standard deviation of 18.91

**Supplementary Note 3** In order to test our assumption that running the base callers in their suggested parameter settings is sound we performed an experiment with XHMM which is the best performing method in our benchmarks. We ran it in also conservative and liberal settings in addition to the suggested setting. The parameter values that correspond to these settings are given in the table below.

| XHMM | minTarget Size | maxTarget Size | minMean TargetRD | maxMean TargetRD | minMean SampleRD | maxMean SampleRD | maxSd SampleRD |
|---|---|---|---|---|---|---|---|
| Conservative | 10 | 1000 | 10 | 5000 | 25 | 2000 | 1500 |
| Suggested | 5 | 10000 | 5 | 5000 | 5 | 2000 | 1500 |
| Liberal | 0 | 100000 | 0 | 50000 | 0 | 20000 | 15000 |

The precision values before and after polishing with DECoNT are given in the table below.

| XHMM | Dup Precision (Unpolished-Polished) | Del Precision (Unpolished-Polished) | Overall Precision (Unpolished-Polished) |
|---|---|---|---|
| Conservative | 0.4758 - 0.6548 | 0.4572 - 0.7120 | 0.4665 - 0.6834 |
| Suggested | 0.4541 - 0.6451 | 0.4144 - 0.7046 | 0.4348 - 0.6704 |
| Liberal | 0.3785 - 0.5543 | 0.3028 - 0.5921 | 0.3406 - 0.5732 |

We observe that the liberal setting results in a worse polished precision $\sim 10\%$. Conservative and suggested setting results are similar. The improvement in precision values are stable across all runs. Thus, we suggest using the default parameter settings for the base callers unless they return insufficient number of calls which prohibit DECoNT training. Then, the parameter choices can be relaxed.

**Supplementary Note 4** The 28 sample taken from 1000 Genomes dataset that were used in CNLearn analysis is given below:

– NA11832, NA12249, NA18968, NA19144
